## Supplementary Data and Figures for "A histomorphological atlas of resected mesothelioma discovered by self-supervised learning from 3446 whole-slide images"

| Option 1 - Epithelioid tumour | | | | | | | |
|---|---|---|---|---|---|---|---|
| | Topic | Options | | | | | |
| 1 | Predominant architectural growth pattern | Tubular | Papillary | Trabecular | Adenomaotoid | Solid | Micropapillary |
| 2 | Second most predominant pattern | Tubular | Papillary | Trabecular | Adenomaotoid | Solid | Micropapillary |
| 3 | Nuclear atypia | Mild | Moderate | Severe | | | |
| 4 | Inflammation* | None-sparse | Mild-moderate | Marked | | | |
| 5 | Necrosis** | None | Some | Universal | | | |
| 6 | Stroma:Tumour ratio | More stroma | Roughly equal | More tumour | | | |
| 7 | Stromal cellularity*** | Low | Moderate | High | | | |
| 8 | ˆBiphasic features? | No | Yes | | | | |
| 9 | Other notable features or comments | free text/annotations | | | | | |
| | ˆBy biphasic, we refer to mixed epithelioid and spindled morphology within individual tiles rather than within the cluster | | | | | | |

| Option 2 - Spindle cells / extracellular matrix | | | | |
|---|---|---|---|---|
| | Topic | Options | | |
| 1 | Cellularity | Low | Medium | High |
| 2 | Architecture | Orderly (parallel) | Less orderly (eg wavy) | Disorderly (random/storiform) |
| 3 | Presence of desmoplastic sarcomatoid morphology | Present | Absent | |
| 4 | Nuclear atypia | None/mild | Moderate | Severe |
| 5 | Inflammation* | None-sparse | Mild-moderate | Marked |
| 6 | Necrosis** | None | Some | Universal |
| 7 | ˆBiphasic features? | No | Yes | |
| 8 | Other notable features or comments | free text/annotations | | |
| | ˆBy biphasic, we refer to mixed epithelioid and spindled morphology within individual tiles rather than within the cluster | | | |

| Option 3 - Non-tumour | | | | | | | | |
|---|---|---|---|---|---|---|---|---|
| Topic | Options | | | | | | | |
| Tiles contain mostly (by area) | Normal/near-normal lung | | | Collagenosis | Haemorrhage | Vessels (wall or lumen) | Airway | Cartilage |
| Second most common feature (by area) | Normal/near-normal lung | Non-neoplastic pathological lung | Elastosis | Collagenosis | Haemorrhage | Vessels (wall or lumen) | Airway | Cartilage |
| Inflammation* | None-sparse | Mild-moderate | Marked | | | | | |
| Necrosis** | None | Some | Universal | | | | | |
| Other notable features or comments | free text/annotations | | | | | | | |
| Inflammation* | None-sparse | Mild-moderate | Marked | | | | | |

**Supplementary Table 1:** Criteria for Pathologist Annotation Sheets

| Leiden Resolution | Dataset | Balanced Accuracy | AUC | Sensitivity | Specificity |
|---|---|---|---|---|---|
| 2.0 | LATTICe-M | 0.79 ± 0.05 | 0.88 ± 0.04 | 0.81 ± 0.1 | 0.77 ± 0.09 |
| 4.0 | | 0.79 ± 0.04 | 0.89 ± 0.03 | 0.81 ± 0.1 | 0.77 ± 0.06 |
| 7.0 | | 0.79 ± 0.04 | 0.88 ± 0.04 | 0.78 ± 0.06 | 0.79 ± 0.08 |
| CLAM | | 0.79 ± 0.03 | 0.87 ± 0.04 | 0.77 ± 0.09 | 0.80±0.05 |
| 2.0 | TCGA-MESO | 0.71 ± 0.03 | 0.8 ± 0.03 | 0.63 ± 0.06 | 0.79 ± 0.04 |
| 4.0 | | 0.78 ± 0.05 | 0.85 ± 0.02 | 0.7 ± 0.05 | 0.85 ± 0.06 |
| 7.0 | | 0.78 ± 0.05 | 0.85 ± 0.04 | 0.67 ± 0.08 | 0.89 ± 0.04 |
| CLAM | | 0.67 ± 0.05 | 0.74 ± 0.01 | 0.55 ± 0.04 | 0.79 ± 0.03 |

**Supplementary Table 2: Subtype classification performance metrics across clustering resolutions and datasets.** Balanced accuracy, area under the curve (AUC), sensitivity, and specificity scores for mesothelioma subtype classification using HPL (across different Leiden clustering resolutions) and CLAM methods on LATTICe-M and TCGA-MESO cohorts. Scores represent means from 5-fold cross-validation following data preprocessing with the edited nearest neighbour (ENN) sampling method in HPL.

| | HPCs | | | Clinical and HPCs | |
|---|---|---|---|---|---|
| Leiden Resolutions | Train | Test | TCGA | Train | Test |
| 2.0 | 0.67 ± 0.0 | 0.65 ± 0.03 | 0.65 ± 0.01 | 0.68 ± 0.01 | 0.66 ± 0.03 |
| 4.0 | 0.69 ± 0.0 | 0.66 ± 0.03 | 0.66 ± 0.02 | 0.7 ± 0.0 | 0.66 ± 0.03 |
| 7.0 | 0.7 ± 0.01 | 0.65 ± 0.04 | 0.66 ± 0.01 | 0.71 ± 0.01 | 0.66 ± 0.04 |
| | Risk Score | | | Clinical and Risk Score | |
| CLAM | 0.60±0.01 | 0.60±0.04 | 0.61±0.0 | 0.62±0.02 | 0.62±0.05 |

**Supplementary Table 3: Concordance indices for patient outcome prediction across datasets and clustering resolutions.** Patient outcome prediction was performed using Cox proportional hazards models, with concordance indices reported across training, testing (LATTICe-M), and external validation (TCGA-MESO) datasets at different Leiden clustering resolutions using 5-fold cross-validation. For the primary dataset, clinical variables (tumour type, TNM stage, and age) were integrated with patient vectors derived from HPL. Additionally, survival analysis was conducted independently using the CLAM multiple instance learning approach, where risk scores were generated through gated attention mechanisms.

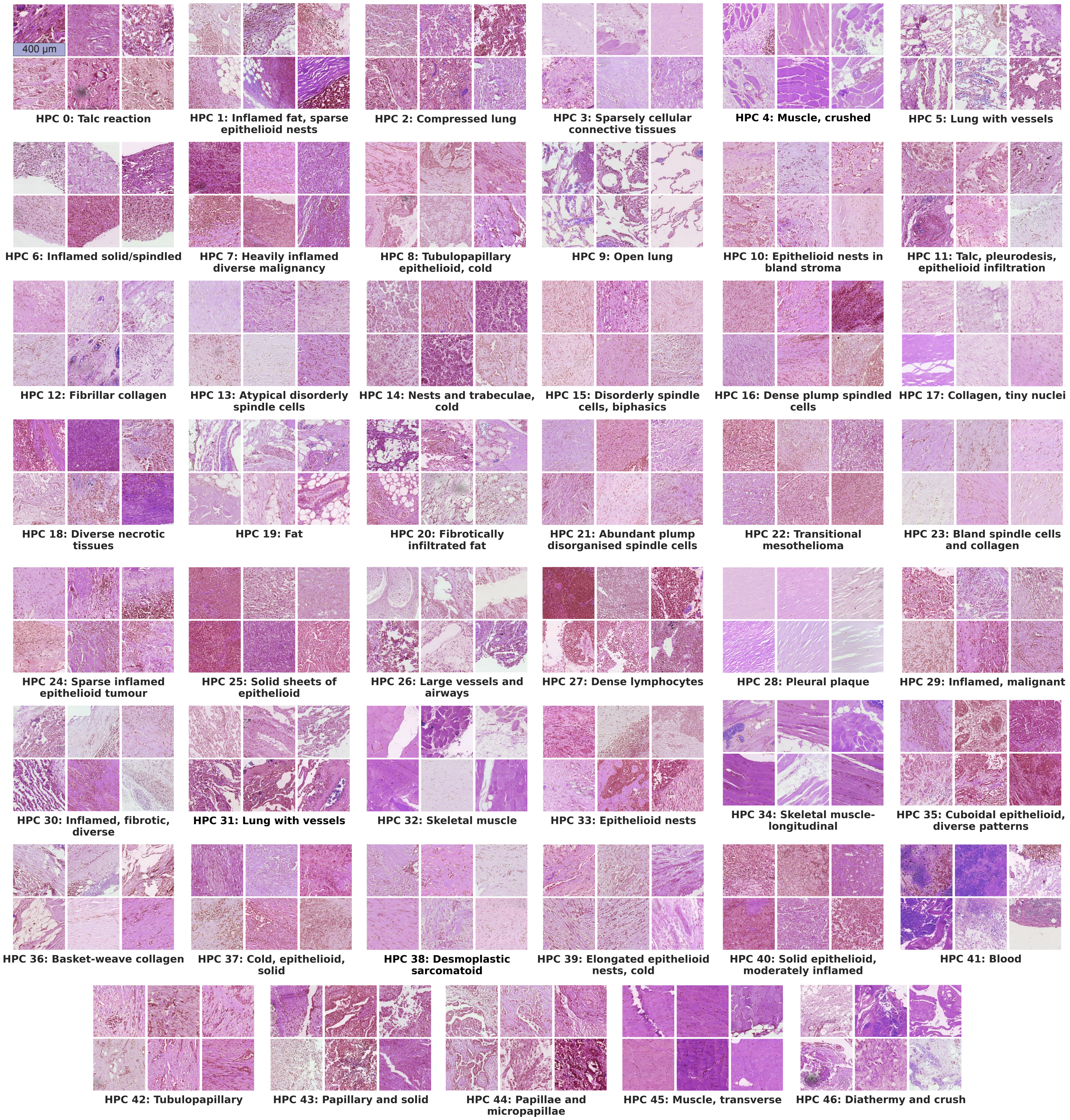


**Supplementary Figure 1: Representative of all HPC tiles in the LATTICe-M dataset.** Examples of tiles from LATTICe-M dataset assigned to all HPCs with short annotations provided from expert pathologists (scale bar, 400 µm).

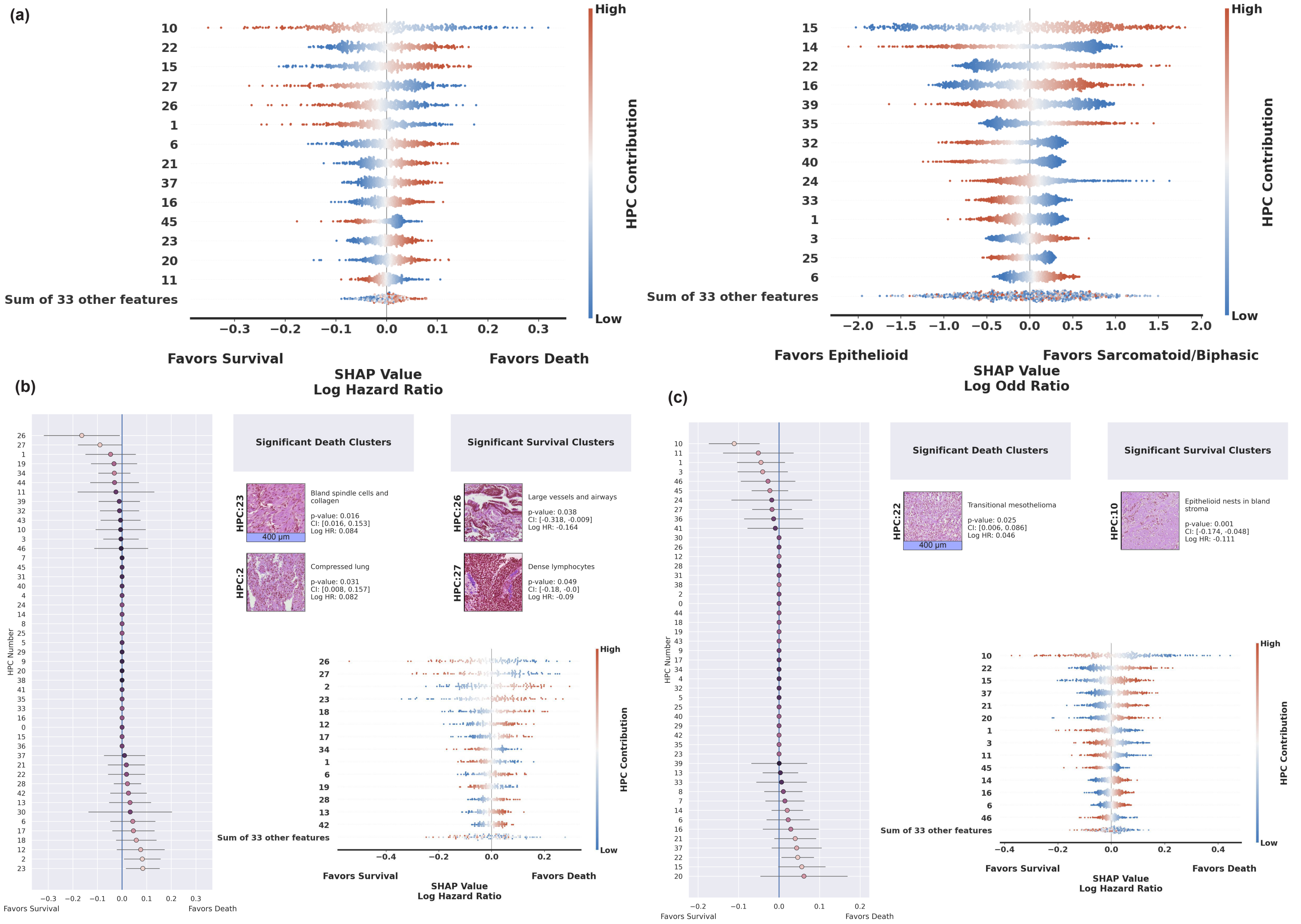


**Supplementary Figure 2: SHAP analysis and subtype-specific Cox model results a** SHAP plots showing the impact of individual HPCs on model predictions for overall survival (left - n= 512) and subtype classification (right - n=3446), ordered by mean absolute SHAP value. **b,c** SHAP summary plots and forest plots from Cox models trained separately on Epithelioid and Sarcomatoid/Biphasic group cases, highlighting subtype-specific prognostic HPCs. All SHAP analyses were performed on case-independent test sets. SHAP values represent each feature's impact on the model's output. Forest plots show log hazard ratios (center) with 95% confidence intervals (error bars), from subtype-specific Cox models. The plots are organized with Sarcomatoid/Biphasic on the left and Epithelioid on the right (scale bar, 400 µm).

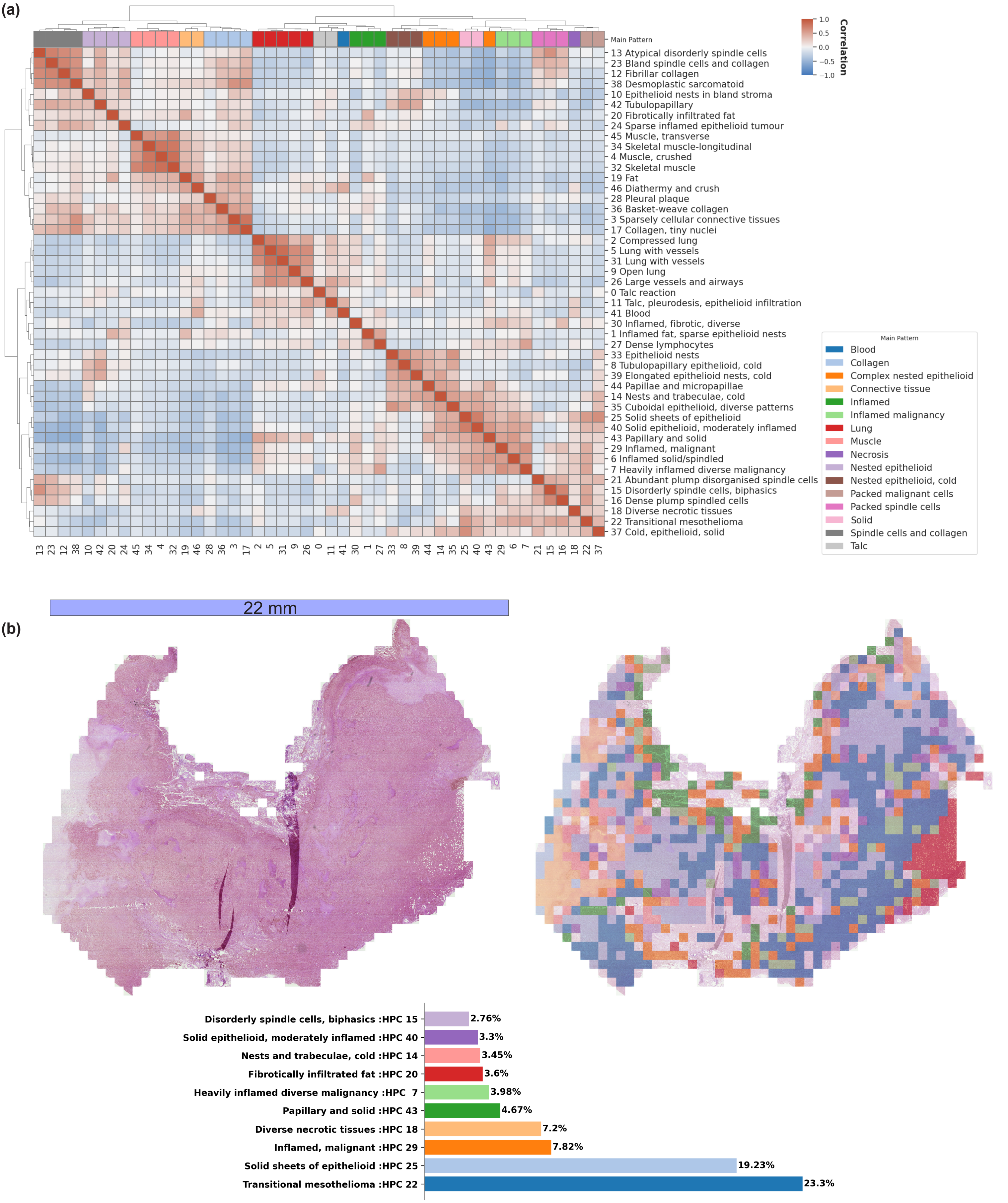


**Supplementary Figure 3: HPC correlation analysis and epithelioid case example.** **a** HPC Pearson (two-sided) correlation heatmap; annotations were independently provided by a pathologist after examination of representative sets of tile images (100 random tiles per HPC). **b** An epithelioid sample case characterised mostly by transitional mesothelioma HPC (scale bar, 22 mm).
